# Central carbon metabolism is an intrinsic factor for optimal replication of a norovirus

**DOI:** 10.1101/434019

**Authors:** Karla D. Passalacqua, Jia Lu, Ian Goodfellow, Abimbola O. Kolawole, Jacob R. Arche, Robert J. Maddox, Mary X.D. O’Riordan, Christiane E. Wobus

**Affiliations:** Department of Microbiology and Immunology, University of Michigan, Ann Arbor, Michigan, USA; Division of Virology, Department of Pathology, University of Cambridge, Cambridge, UK.

**Keywords:** Caliciviridae, norovirus, metabolism, glycolysis, oxidative phosphorylation, pentose phosphate pathway

## Abstract

The metabolic pathways of central carbon metabolism, glycolysis and oxidative phosphorylation (OXPHOS), are important host factors that determine the outcome of viral infections and can therefore be manipulated by some viruses to favor infection. However, mechanisms of metabolic modulation and their effects on viral replication vary widely. Herein, we present the first metabolomics profile of norovirus-infected cells, which revealed increases in glycolysis, OXPHOS, and the pentose phosphate pathway (PPP) during murine norovirus infection. Inhibiting glycolysis with 2-deoxyglucose (2DG) in transformed and primary macrophages revealed that host cell metabolism is an important factor for optimal murine norovirus (MNV) infection. 2DG affected an early stage in the viral life cycle after viral uptake and capsid uncoating, leading to decreased levels of viral protein translation and viral RNA replication. The requirement of central carbon metabolism was specific for MNV (but not astrovirus) infection, independent of the Type I interferon antiviral response, and unlikely to be due to a lack of host cell nucleotide synthesis. MNV infection increased activation of the protein kinase Akt, but not AMPK, two master regulators of cellular metabolism, suggesting Akt signaling may play a role in upregulating central carbon metabolism during norovirus infection. In conclusion, our findings suggest that the metabolic state of target cells is an intrinsic host factor that determines the extent of norovirus replication and implicates metabolism as a virulence determinant. They further implicate cellular metabolism as a novel therapeutic target for norovirus infections and improvements of current human norovirus culture systems.

**IMPORTANCE:** Viruses depend on the host cells they infect to provide the machinery and substrates for replication. Host cells are highly dynamic systems that can alter their intracellular environment and metabolic behavior, which may be helpful or inhibitory for an infecting virus. In this study, we show that macrophages, a target cell of murine norovirus (MNV), increase central carbon metabolism upon viral infection, which is important for early steps in MNV infection. Human noroviruses (hNoV) are a major cause of gastroenteritis globally, causing enormous morbidity and economic burden. Currently, no effective antivirals or vaccines exist for hNoV, mainly due to the lack of high efficiency *in vitro* culture models for their study. Thus, insights gained from the MNV model may reveal aspects of host cell metabolism that can be targeted for improving hNoV cell culture systems and for developing effective antiviral therapies.

## INTRODUCTION

Viruses are obligate intracellular parasites. Thus, their biology is entirely dependent on the physiology of the host cells they infect. One increasingly appreciated aspect of virus-host interaction is cellular metabolism (1–4). Historically, cellular metabolism has been considered mainly in terms of its role in cellular energy homeostasis. However, metabolism and metabolic “cross talk” are increasingly being appreciated as crucial aspects in a range of cellular processes such as proliferation and cell death (5), the activation and functioning of the immune system (6, 7), autophagy (8, 9) and in the establishment of infectious disease (10). Indeed, a wide range of pathogens including parasites (11), bacteria (12–14) and viruses (3) have been shown to affect and to be affected by their hosts’ metabolic activity. Of note, the controlled modulation of metabolism in immune cells has been shown to be a key feature in adaptive and innate immune responses (6, 14–16), and these findings have given rise to an entire field referred to as “immunometabolism” (17–20). For example, macrophages adapt to a variety of metabolic profiles depending upon the specific signals they sense (21, 22). Specifically sensing through different Toll-Like Receptors (TLR) in myeloid cells can initiate any combination of up- and/or down-regulation of glycolysis and OXPHOS (23). Thus, metabolic processes are a vital feature of the immune system for effectively combating viral infections, or an Achille’s heel of the host cell that can be manipulated by invading pathogens for their own advantage.

Eukaryotic cellular metabolism encompasses a wide range of catabolic and anabolic processes, and various aspects of host metabolism have been linked to viral infections. In particular, the major pathways of central carbon metabolism, glycolysis and oxidative phosphorylation (OXPHOS), have been investigated for their role in viral infection. For example, Kaposi’s sarcoma herpesvirus (KSHV) suppresses aerobic glycolysis and OXPHOS to foster cellular, and thus viral, survival (24). In contrast, an array of diverse viruses such as herpes simplex virus 1, HIV-1, rubella virus, white-spot syndrome virus, dengue virus, rhinovirus, hepatitis C virus, influenza virus, and adenovirus (25–33) have been shown to initiate a host cell response characterized by an increase in glycolysis, resulting in a more hospitable intracellular environment for viral replication. However, the specific ways in which viral infections initiate metabolic responses, and how these responses affect viral infection, vary substantially. Disentangling the unique metabolic responses of host cells upon viral infection, especially in regard to glycolysis and OXPHOS, may help in the development of broadly acting antiviral therapies.

Human noroviruses (hNoV) are non-enveloped, positive-sense, single-stranded RNA viruses of the *Caliciviridae* family that cause the majority of acute non-bacterial gastroenteritis globally (34–37). In addition to the public health burden, the economic burden of hNoV infections is enormous, with global costs estimated at $60 billion (US$) annually (35, 36). Currently, there are no licensed vaccines or antivirals that are effective against hNoV infections. Although advances have been made in developing *in vitro* model systems for studying hNoV (38–42), the field still lacks a highly efficient, easy-to-use cell culture model. Therefore, murine norovirus (MNV) remains a powerful tool for investigating general norovirus biology (43–45). The goal in the current study was to identify aspects of host cell metabolism that are important for modulating MNV replication. Such findings may enable the development of more efficient hNoV culture systems and/or antiviral therapies and vaccines for hNoV in the future (46).

With these goals in mind, we performed the first metabolomic analysis of norovirus infection. Our analysis demonstrated that MNV infection of macrophages causes changes in the host cell metabolic profile characterized by an increase in central carbon metabolism. Inhibition of glycolysis with 2-deoxyglucose (2DG) severely attenuated MNV, but not human astrovirus VA1, infection *in vitro*. Inhibition occurred at the level of replication, as we observed a lag in the appearance of viral proteins in infected cells with a concomitant lag in viral genome replication, but no effect on viral uptake or uncoating. Inhibition of MNV infection by 2DG was not rescued by addition of nucleotides and was independent of Type I interferon responses. Investigations of the two master regulators of cellular metabolism, Akt and AMPK, revealed that MNV infection caused an increase in Akt activation, while inhibition of Akt signaling reduced both cellular glycolysis and MNV infection. Overall, our findings identify central carbon metabolism as an intrinsic host factor important for optimal MNV infection of macrophages. Since noroviruses have a tropism for immune cells (47) and specific immune cell subsets are characterized by different metabolic profiles (48, 49), these findings may have implications for viral pathogenesis and the development of improved hNoV culture systems.

## RESULTS

### Targeted metabolomics survey identifies multiple metabolites that increase during MNV-1 infection in RAW 264.7 cells

Viral infections can cause changes in host cell metabolism that are important for viral replication (1, 3). In our efforts to identify host cell factors that are important for successful norovirus (NoV) infection, we hypothesized that infection of macrophages with murine norovirus (MNV) causes changes in central carbon metabolism of host cells that are beneficial or required for optimal viral infection. MNV-1 (CW3 isolate) is an acute strain of murine norovirus that has a natural tropism for macrophages *in vivo* and is particularly efficient at infecting transformed murine macrophages RAW 264.7 (RAW) (44). Thus, we performed a targeted metabolomics profiling of MNV-infected RAW cells to identify changes in the amount of host cell metabolites from glycolysis, the tricarboxylic acid cycle (TCA) and others.

A targeted mass spec analysis of metabolites isolated from MNV-1 infected RAW cells (MOI=5) after eight hours of infection (approximately one replication cycle) revealed multiple metabolites that were significantly increased in infected cells compared to mock cells, or unchanged, but no metabolites that were significantly decreased during infection (**Fig. 1**) (**Supplemental Tables 1 and 2**). In particular, an increase in select metabolites from glycolysis (fructose-bisphosphate, 2- and 3-phosphoglycerate, dihydroxyacetone-phosphate), the pentose phosphate pathway (PPP) (6-phosphogluconate) and the TCA cycle (citrate/isocitrate, malate) suggest that both energy generating pathways of glycolysis and oxidative phosphorylation are increased during MNV infection (**Fig. 1A**). Notably, overall levels of ATP were higher in infected cells compared to mock (**Fig. 1A**), indicating an overall increase in RAW cell metabolism as a result of viral infection. The detection of a significant increase in metabolites in cell culture is particularly noteworthy, since MNV-infected cultures represent a heterogeneous population of infected and uninfected cells, since not all cells get infected by MNV even when experiments are done with a high MOI (50).

**Fig 1.**
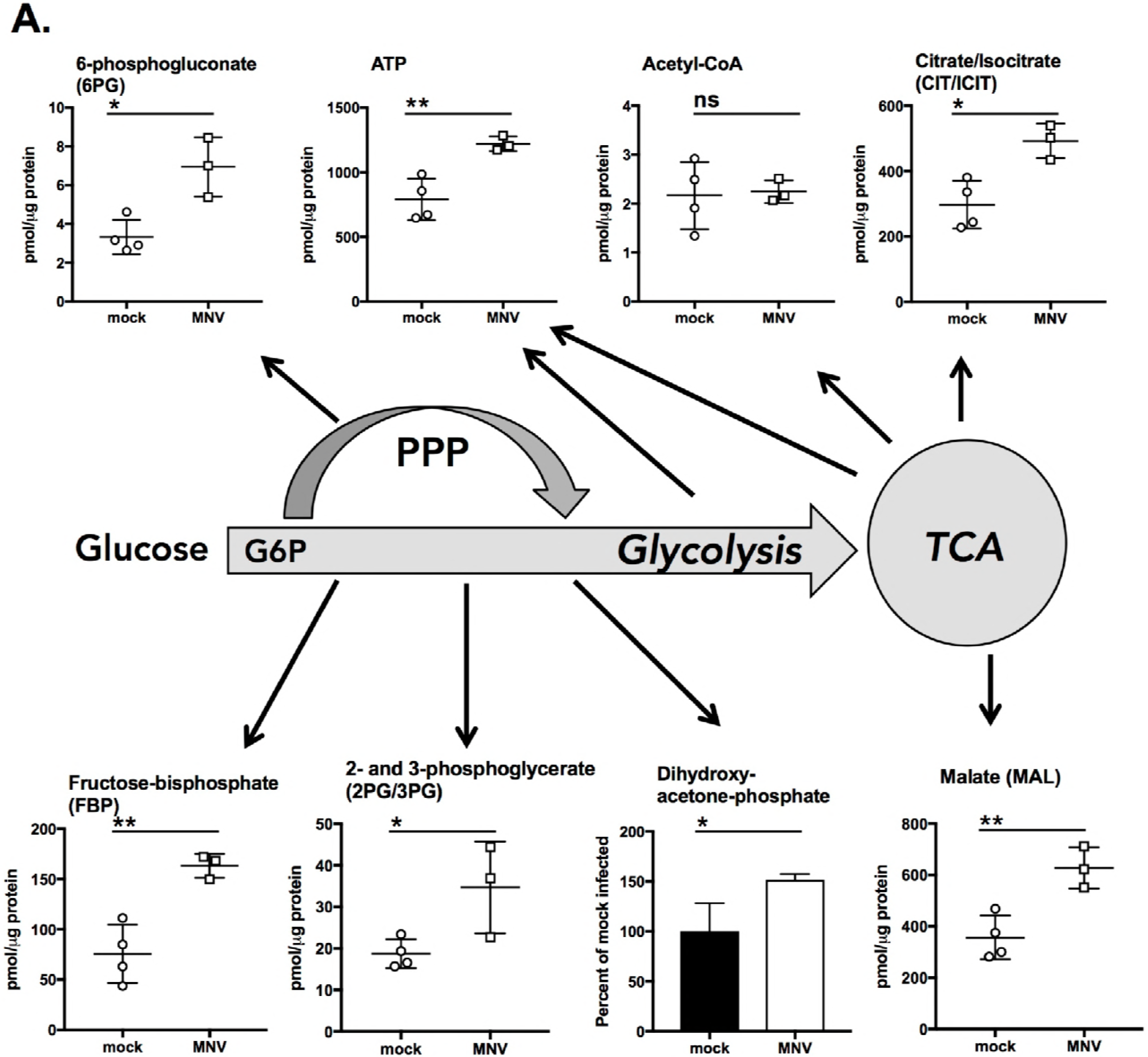
Metabolomics survey of RAW 264.7 cells infected with MNV-1 reveal several metabolic pathways that are increased during infection. (A) Measurements of select metabolites from central carbon metabolism, including glycolysis, the Pentose Phosphate Pathway (PPP), and the Tricarboxylic Acid Cycle (TCA). (B) Metabolites from Xanthine biosynthesis (Purine metabolism), and (C) the UDP-Glucuronate pathway (Glucuronic acid pathway). Schematics of the metabolic pathways shown are simplified for clarity. All metabolites assayed are listed in Tables 1 and 2 with mean and standard deviation for three MNV-1 infected samples (MOI = 5) and four mock-infected samples (mock cell lysate). Infection was 8 hours. ♦Indicates where in the pathway UTP is consumed. Analyses performed in Metaboanalyst using student’s t-test. \**P*<0.05; \*\**P*<0.01; ns = not significant.

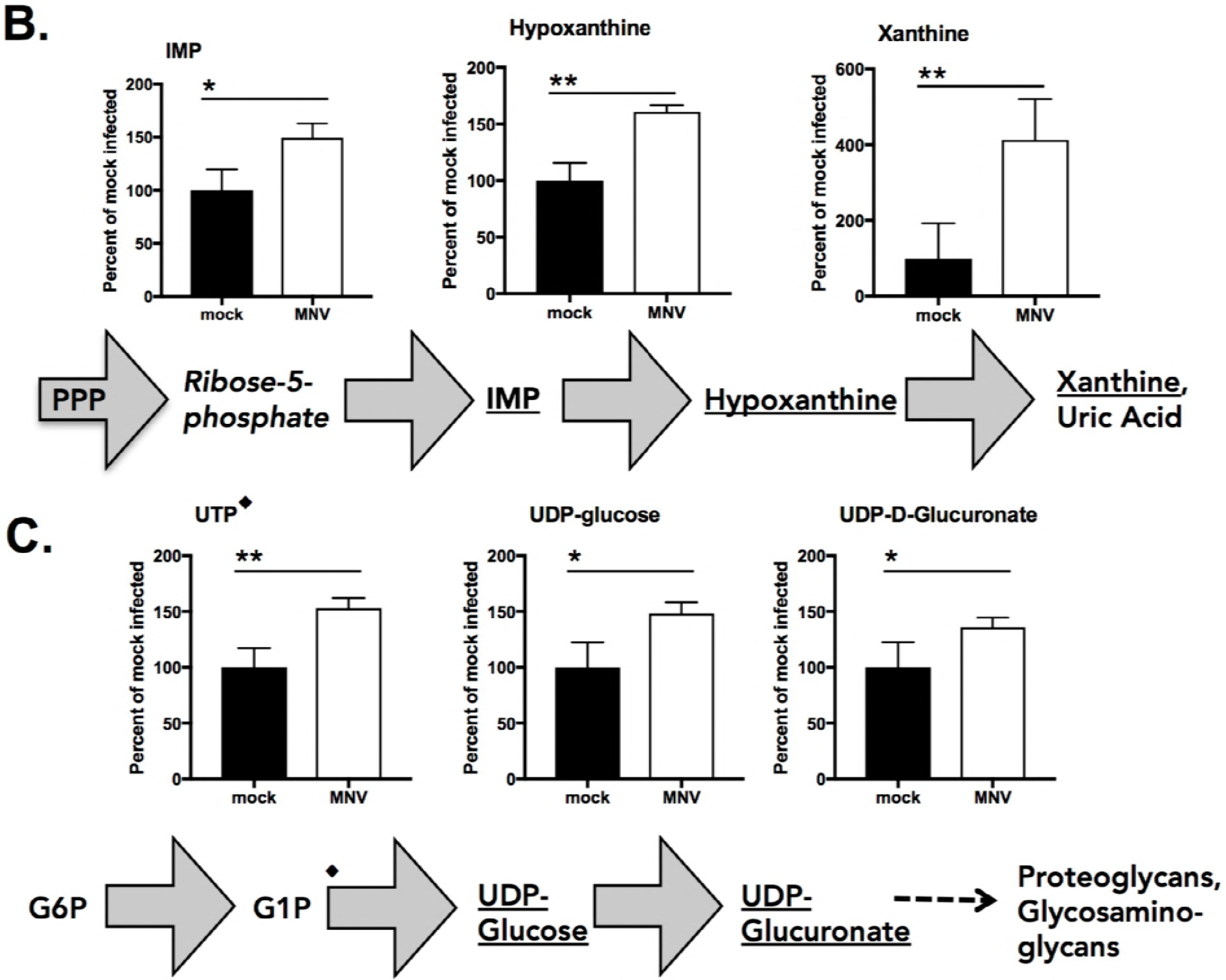

Another group of metabolites that increased in RAW cells during MNV infection include inosine-monophosphate (IMP), hypoxanthine and xanthine (**Fig. 1B**). These metabolites are part of a pathway involved in adenosine catabolism that can result in the production of uric acid, a potent immune signal (16), and potentially reactive oxygen intermediates, which can have signaling and antimicrobial activity. Upregulation of enzymatic activity in this pathway and an increase in the resulting metabolites has been observed in the liver of mice infected with several RNA and DNA viruses (51), in the lungs and tissues of influenza virus-infected mice (52), and in mice infected with rhinovirus (53), and thus may represent a generalized cellular response to viral infection.

Lastly, the metabolites uridine tri-phosphate (UTP), UDP-glucose, and UDP-D-glucuronate were also increased in MNV-infected RAW cells (**Fig. 1C**). These metabolites are part of the glucuronic acid pathway that can lead to the generation of proteoglycans and other glycosylated forms of proteins (54) that have variable roles, including as potential extracellular signals (55, 56). Indeed, many hNoV strains, including the clinically relevant genogroup II, genotype 4 viruses, are able to bind to host extracellular glycans, i.e., histo-blood group antigens (57, 58). Collectively, our metabolomics survey suggests that macrophages respond to MNV infection by increasing: (i) the energy- and metabolite-generating pathways glycolysis and oxidative phosphorylation; (ii) adenosine catabolism, which may be a part of the general innate immune response; and, (iii) the glucuronic acid pathway, which may have effects on cellular protein glycosylation.

### 2DG reduces MNV-1 infection in RAW cells and bone marrow-derived macrophages

Metabolomics profiling of MNV infected RAW cells suggested that glycolysis and OXPHOS are increased during viral infection. But whether this increase creates an intracellular environment more supportive for viral replication, or rather represents an anti-viral immune strategy of the host cell, is unclear from such a survey. To test whether host cell glycolysis is supportive for effective MNV infection of macrophages in generating building blocks, viral infection was measured *in vitro* in the presence of the potent and commonly used glycolysis inhibitor 2-Deoxyglucose (2DG), a glucose analog that blocks early glycolysis (59, 60).

RAW cells were infected with MNV-1 at an MOI of five for one hour. Medium containing 10 mM 2DG was then added post-infection to exclude direct effects of the compound on virions. After an eight-hour incubation (one viral replication cycle), a greater than two log_10_ decrease in the number of infectious viral particles in 2DG-treated cells was observed by plaque assay (**Fig. 2A**). RAW cells are a transformed cell line and generally engage in active “Warburg-effect” glycolysis (61). We therefore repeated the experiment in primary bone marrow-derived macrophages (BMDM) isolated from Balb/c mice to determine whether glycolysis is also relevant in non-transformed cells. 2DG treatment of BMDM caused an average one log_10_ decrease in viral loads after eight hours (**Fig. 2B**). 2DG treatment did not inhibit RAW viability during an eight-hour treatment (**Fig. 2C**), but did reduce RAW cell viability by about 30% after 24 hours (**Fig. S1A**).

**Fig 2.**
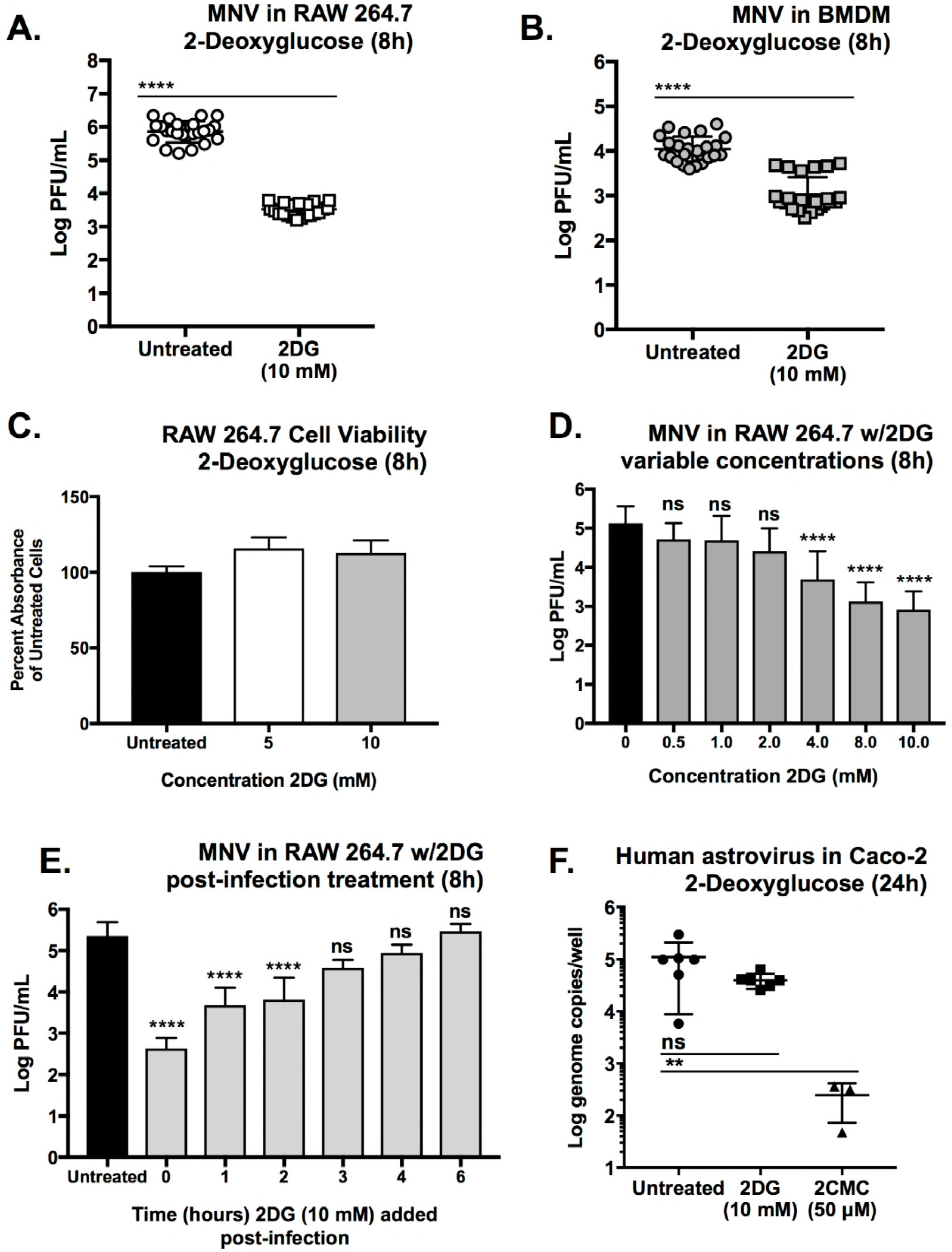
Effects of 2-Deoxyglucose (2DG) on MNV-1 and human astrovirus VA1 infection *in vitro*. (A) 2DG (10 mM) reduces MNV-1 infection in RAW cells (~2 log_10_), and (B) primary Bone-marrow derived macrophages (~1 log_10_) (BMDM-Balb/C mice). 8-hour infections with MOI=5. (C) Cell viability assay (Resazurin reagent) showing that 2DG does not reduce RAW cell viability during 8-hours of exposure. Cell viability at 24 hours in Figure S1A. (D) Effects of different concentrations of 2DG on MNV-1 infection in RAW cells. (E) MNV-1 infection in RAW cells with 2DG added at different times post-infection. (F) 2DG does not affect infection of human astrovirus VA1 in Caco-2 cells (2DG 10 mM; 2CMC positive control 50 μM). Toxicity of 2DG on Caco-2 cells in Fig S1E. (A, B, D and E) measured by Plaque Assay. Astrovirus in (F) was measured by RT-qPCR of viral RNA. Mann-Whitney test used for (A, B, F) where \*\*\*\**P*<0.0001. Kruskal-Wallis test with Dunn’s multiple comparisons test used for (E) where \*\*\*\**P*<0.0001 and ns = not significant. Experiments represent combined data from at least three independent experiments except (F), which represents two experiments.

Since RAW cells were grown in medium replete with glucose (~25 mM), putatively creating a competitive metabolic situation between glucose and 2DG, we next determined the minimal concentration of 2DG that significantly inhibited viral infection in RAW cells. Findings from a dose-response study performed in the presence of glucose demonstrated that 2DG inhibited MNV-1 infection in a dose-dependent manner with the lowest significant inhibition at 4.0 mM (**Fig. 2D**).

To determine the point during the infectious cycle that 2DG exerts its inhibitory effect on MNV, a time-of-addition study was performed. RAW cells were infected with MNV-1 and 2DG added to the medium at variable times post-infection. The results showed that 2DG had a significant effect on MNV infection when added to the culture up to two hours post-infection (**Fig. 2E**), suggesting that glycolysis is important for early steps in the viral life cycle.

These data are consistent with the notion that glycolysis is providing necessary building blocks for viral replication. Thus, to determine whether 2DG might be exerting a generalized anti-viral response in any transformed cell line against any virus, we tested viral infection of a different ssRNA virus, human astrovirus VA1, which is readily propagated in Caco-2 cells (62). Surprisingly, 2DG did not significantly inhibit human astrovirus infection *in vitro* (**Fig. 2F**), suggesting that the MNV phenotype in RAW cells and in BMDM is specific to MNV.

Taken together, these data demonstrate that host cell glycolysis, whether in primary or transformed cells, contributes to optimal MNV infection in macrophages. They further suggest that glycolysis is an intrinsic host factor that modulates infection in a virus-specific manner.

### 2DG treatment inhibits MNV-1 (-) strand vRNA and viral non-structural protein production

Post-infection treatment of RAW cells with 2DG suggested that host cell glycolysis is important for early stages of MNV infection (**Fig. 2E**). To more accurately pinpoint the stage in the viral infectious cycle at which glycolysis is important, RAW cells were transfected directly with viral RNA (vRNA) in order to bypass the steps of binding, uptake and virion uncoating. 2DG treatment of transfected RAW cells resulted in about a two log_10_ reduction of infectious virus particle production after 12 hours (**Fig. 3A**) and a one log_10_ reduction at 24 hours (**Fig. S2**), suggesting that 2DG does not affect virion binding or genome uncoating of MNV.

**Fig 3.**
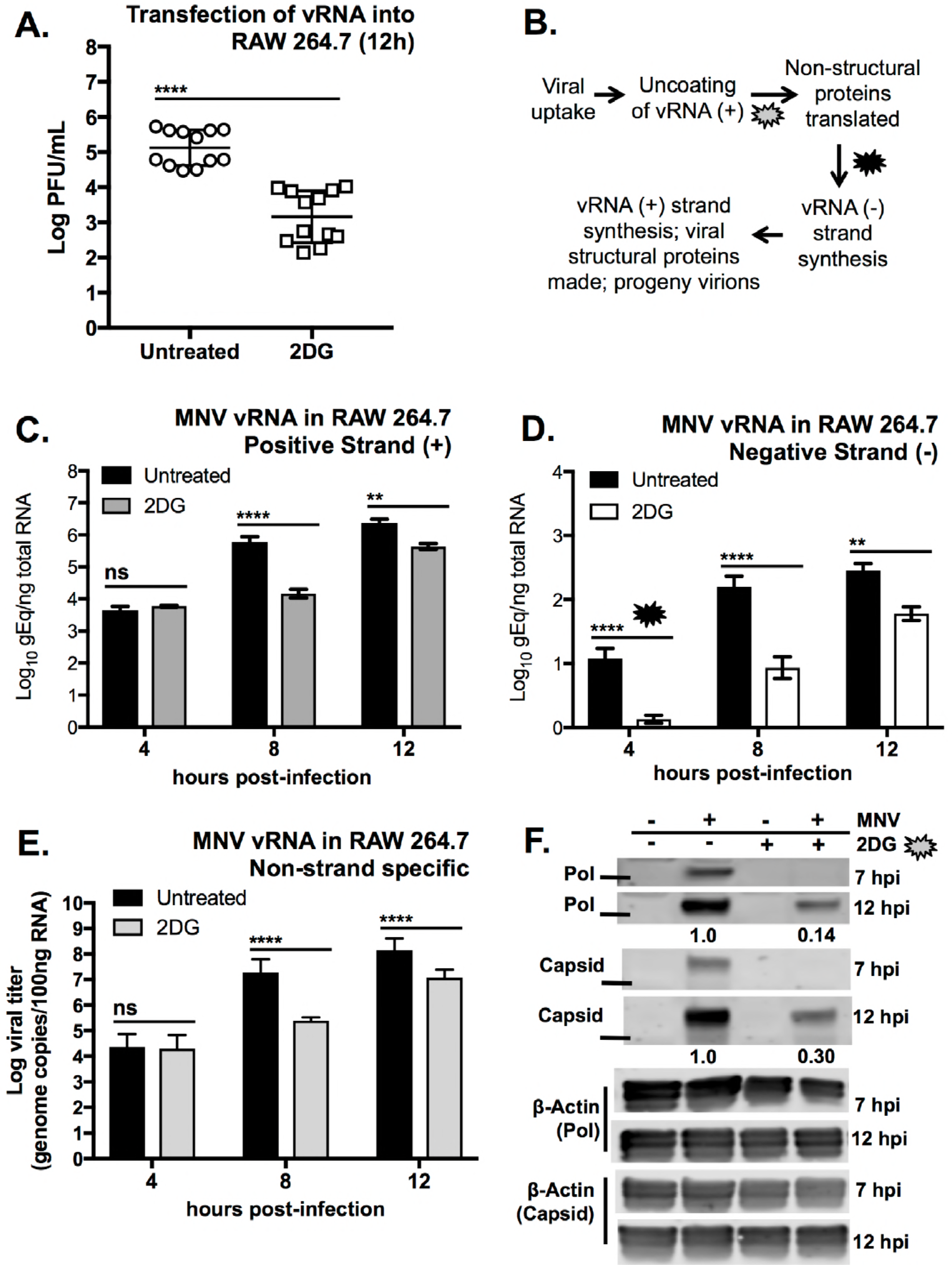
2DG treatment inhibits MNV infection early after viral uptake and uncoating. (A) MNV-1 viral RNA (vRNA) was transfected into RAW cells and then treated with 10 mM 2DG. Data are from two independent experiments. (B) A simplified overview of the events in the MNV-1 life cycle. Callouts indicate points of the viral life cycle that may be affected during 2DG treatment. (C & D) Strand-specific RT-qPCR of (C) plus (+) and (D) minus (-) MNV vRNA strands from RAW cells infected with MNV-1 for 4, 8 and 12 hours with and without 2DG treatment (10 mM). (E) Taq-Man RT-qPCR of total MNV-1 viral RNA (non-strand specific) from the same RNA samples used for (C) and (D). Data are combined from three independent experiments with three replicates per experiment. (F) Western blot analysis of non-structural (Pol) and structural (Capsid) viral proteins after 7 and 12-hour infection of RAW cells in untreated and 2DG treated cells. β-Actin was used as a loading control for overall protein loading content. Solid line indicates 50 kDa ladder. Data shown are a representative Western blots from two independent experiments. Numbers below blots indicate densitometry measurement of protein in 2DG relative to untreated cells at 12 hours (average of two experiments). Mock-infected cells served as negative control. Mann-Whitney test used for (A) where \*\*\*\**P*<0.0001. Two-way ANOVA with Dunnett’s multiple comparisons test used for (C, D and E) where \*\**P*<0.01; \*\*\*\**P*<0.0001; ns = not significant. PFU=plaque forming units.

MNV is a single-stranded, positive (+) strand, non-enveloped virus, and so the viral life-cycle involves uptake of viral particles, uncoating of the (+) strand vRNA, direct translation of the (+)-sense genome to produce the non-structural proteins (including the viral RNA polymerase), followed by viral negative (-) RNA strand synthesis for eventual production of new (+) strand vRNA, structural coat proteins, and progeny virion assembly (**Fig. 3B**). To measure vRNA production during 2DG treatment, we isolated RNA over the course of a 12-hour infection and assessed relative amounts of total and plus- and minus-strand vRNA (**Figs. 3C, D, and E**) (63). At four hours post-infection (hpi), no difference in the quantity of (+) strand and total vRNA was observed (**Figs. 3C and E**), indicating the same amount of virus infected the cells, confirming 2DG has no significant effect on viral binding and entry. However, there is a significant reduction in the amount of (-) strand vRNA at four hpi in 2DG-treated cells (**Fig. 3D**). At 8 and 12 hpi, there is significantly less vRNA overall for all species of RNA assessed (**Fig 3C-E**). These data demonstrated although vRNA replication occurs in 2DG-treated cells, a lag occurred in transcription of (-) strand vRNA.

Since translation of the non-structural proteins precedes (-) strand vRNA synthesis, we next assessed the quantity of MNV non-structural protein using anti-ProPol/NS6&7 and anti-capsid antibodies by Western blot during 2DG treatment. Cells treated with 2DG contained no detectable Pol or VP1 proteins at 7 hpi, while reduced amounts of these proteins were present at 12 hpi (**Fig. 3F**). These data indicate that host cell glycolysis is important for an early step in viral replication after delivery of the viral RNA into the cytosol.

Taken together, inhibition of glycolysis with 2DG did not affect the ability of RAW cells to internalize infectious virions, but it caused a delay in the translation of viral non-structural proteins and negative-strand RNA synthesis. It is currently unclear whether the decrease in (-) strand RNA levels are due to the reduced levels of viral non-structural proteins, including the viral polymerase, or a direct inhibitory effect of 2DG on vRNA synthesis.

### Inhibiting the pentose phosphate pathway reduces MNV infection of RAW cells

The metabolomics survey outlined in Figure 1 demonstrated that the first metabolite produced from Glucose-6-phosphate in the oxidative half of the Pentose Phosphate Pathway (PPP), 6-phosphogluconate, was more abundant in MNV-infected cells. This suggested that the PPP, which branches off glycolysis at the early stage of glucose phosphorylation (64), may also be important for MNV infection in RAW cells. Also, since 2DG interferes at the level of glucose phosphorylation, the viral inhibition caused by 2DG may be due to interference with the PPP.

Therefore, to test the importance of the PPP for MNV infection, we used the inhibitor of the PPP enzyme glucose-6-dehydrogenase, 6-Aminonicotinamide (6AN). Treatment with 500 μM 6AN after MNV-1 infection caused a one log_10_ reduction in the production of infectious MNV-1 after eight hours (**Fig. 4A**). 6AN was non-toxic to RAW cells up to 1.0 mM during eight hours (**Fig. 4B**), whereas all concentrations of 6AN tested caused an approximately 30% reduction in cell viability after 24 hours (**Fig. S1B**).

**Fig 4.**
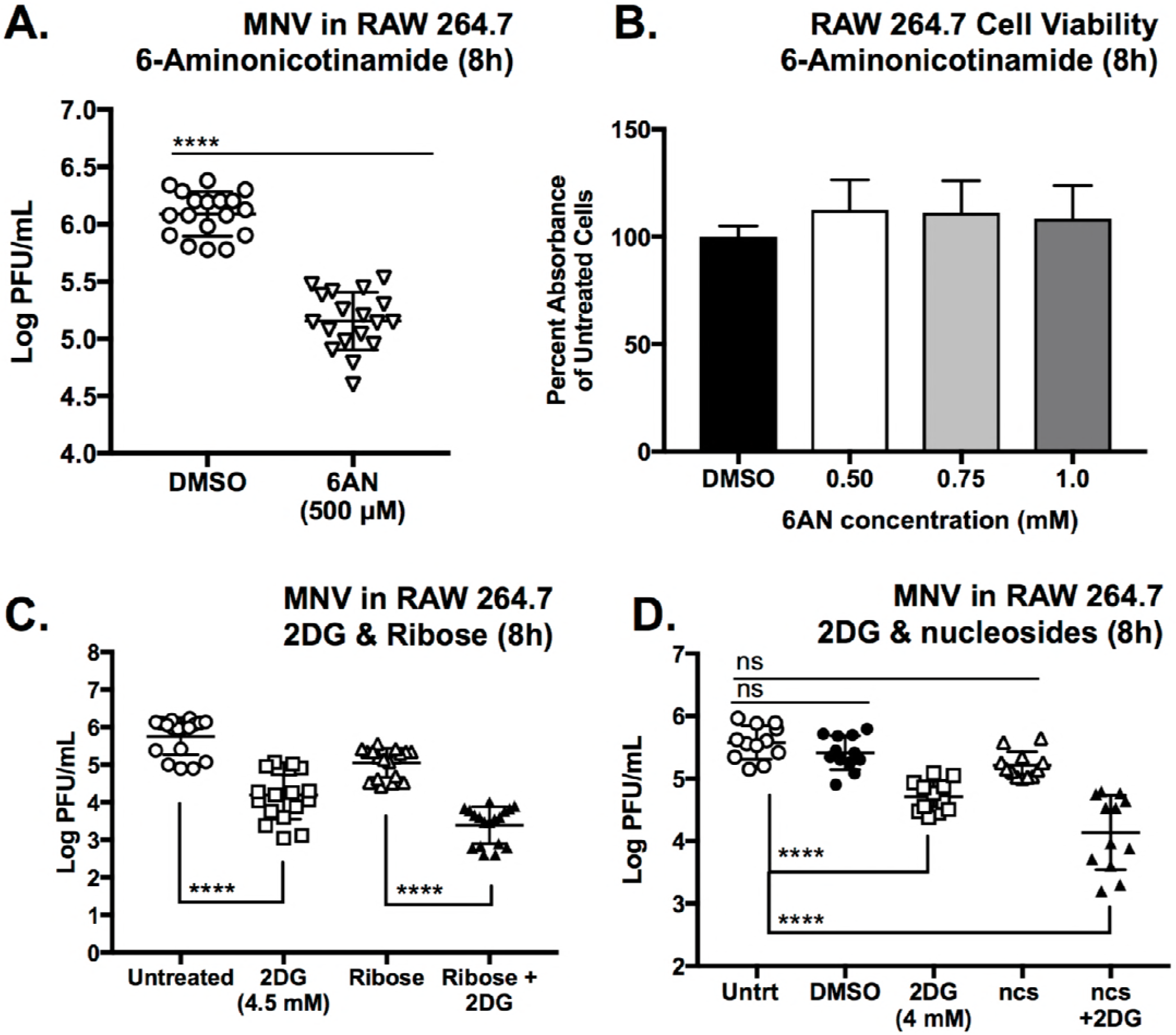
The pentose phosphate pathway makes a minor contribution to MNV infection of RAW cells. (A) 6-Aminonicotinamide (6AN) (500 μM), the inhibitor of 6-phosphogluconate dehydrogenase, reduces MNV infection in RAW cells (MOI = 5). (B) Resazurin cell viability assay of RAW cells treated with indicated concentration of 6AN for 8 hours (see S1B for 24h). (C) Supplementing MNV-infected RAW cells with 50 mM Ribose or (D) 50 μM nucleosides (ncs) does not alleviate the viral growth inhibition caused by 4.5 or 4.0 mM 2DG treatment after 8-hour infection. Nucleosides (ncs) used in (D) were 50 μM each of adenosine, guanosine, thymidine, cytidine, and uridine. RAW cells were treated overnight before infection with nucleosides, and again supplemented with nucleosides after infection with MNV. Mann-Whitney test in (A). Kruskal-Wallis test with Dunn’s multiple comparisons test in (C) and (D). \*\*\*\**P*<0.0001; ns = not significant. PFU=Plaque Forming Units. DMSO is vehicle control used in v/v match to 6AN or ncs treatment. Data represent combination of three independent experiments.

Inhibition of MNV infection by 2DG may be partially due to its effect on the PPP by depleting ribose nucleotides, one of the major end products of the PPP. Alphaviruses, which rely on host cell glycolysis via PI3 kinase signaling, are partially rescued for viral replication with ribose supplementation when PI3 kinase signaling is inhibited (65). Therefore, we infected RAW cells with the minimal amount of 2DG that still causes a significant reduction in MNV infection (4 mM), and supplemented the cultures with ribose alone (**Fig. 4C**) or pre-supplemented cells with a mix of five ribonucleosides (**Fig. 4D**). Neither treatment was sufficient to increase viral titers during 2DG inhibition. These data suggest that, at least under the conditions tested, viral inhibition from 2DG is caused by cellular changes other than nucleotide availability.

### 2DG viral inhibition is independent of the type I interferon response

The mechanism of viral inhibition by 2DG could be due to a variety of cellular perturbations that are caused by a decrease in glycolysis. MNV infection in RAW cells induces a strong innate immune response, including interferon induction (66). Type I Interferons in turn are able to affect host cell metabolism (67, 68), and exhibit a strong anti-MNV response (69–71). Therefore, we determined whether 2DG inhibition of viral replication was dependent on type I interferon responses. Wild-type C57BL/6 BMDM and BMDM lacking the type I interferon receptor (IFNAR1^-/-^) were infected with MNV-1 for one hour and then treated cells with 10 mM 2DG. After eight hours, both WT and IFNAR1^-/-^ cells had reduced viral titers following 2DG treatment compared to untreated cells (**Fig. 5**). These data demonstrate that the inhibition of MNV infection by 2DG is independent of the antiviral type I interferon response.

**Fig 5.**
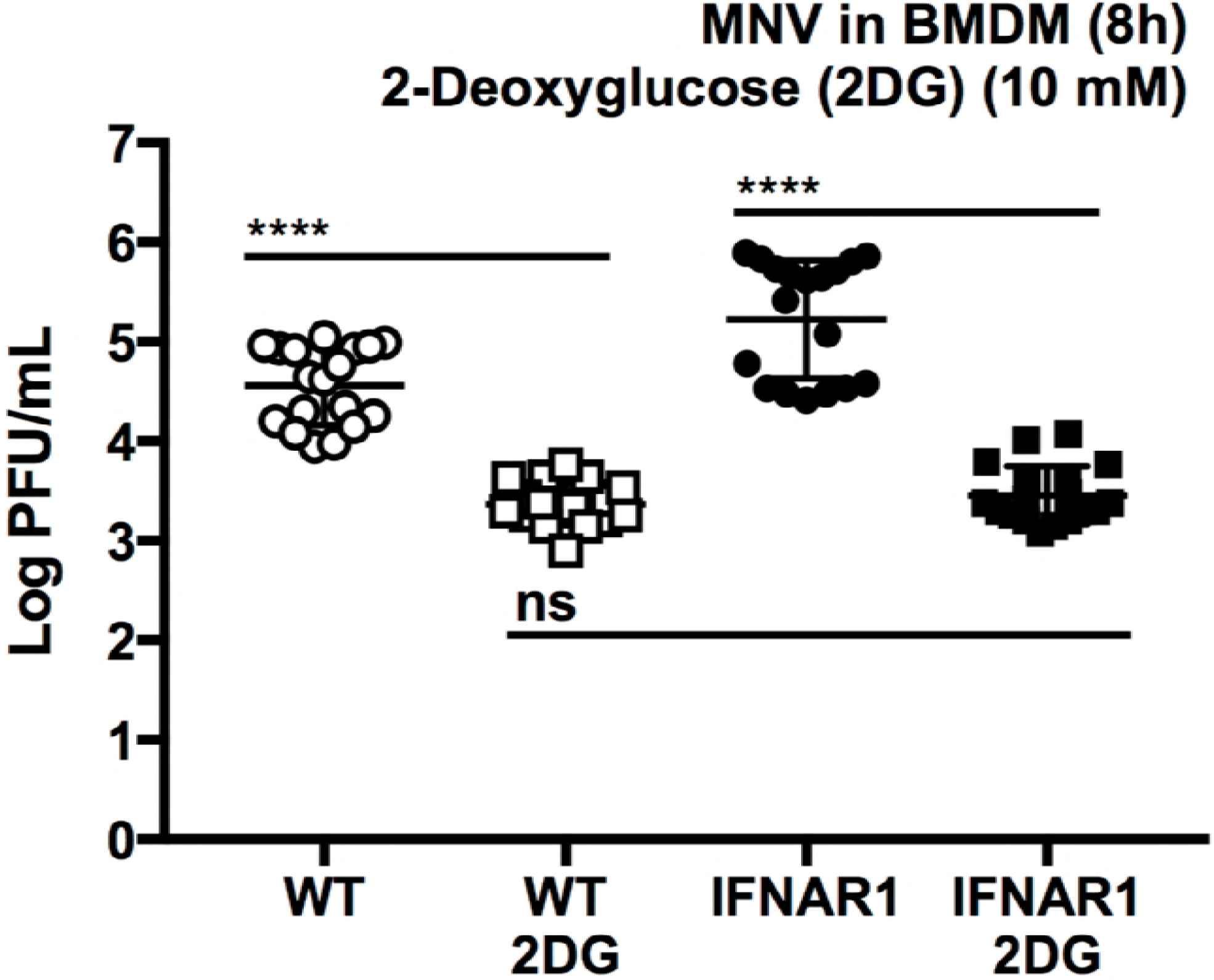
2DG inhibition of MNV infection is independent of the Type I interferon response. 2DG treatment reduces MNV infection in both WT BMDM and in BMDM lacking the Type I Interferon Receptor (IFNAR1 Knockout Cells) (WT-2DG versus IFNAR-2DG). Kruskal-Wallis test with Dunn’s multiple comparisons post-test. \*\*\*\**P*<0.0001; ns = not significant. Data represent a combination of three independent experiments.

### MNV-1 infection increases activation of Akt but not AMPKα

To identify cellular signaling pathways that underlie the observed metabolic changes during MNV infection, we focused on two master regulators of metabolic control in cells, PI3-kinase/Akt and AMPK (72–80). In mammals, AMPK is able to sense the energetic status of cells, specifically the ratio of AMP and ADP relative to ATP, and can promote fatty acid oxidation and the expression of mitochondrial proteins (81–83). Western blot analysis of RAW cells revealed very low levels of total AMPKα protein, and no increases in phosphorylation at Thr172 between mock- and virus-infected cells, or between untreated and 2DG-treated cells were observed (**Fig. 6A and Fig. S3**). These data demonstrate that AMPK is not involved in the energetic changes in RAW cells that we have observed during MNV infection.

**Fig 6.**
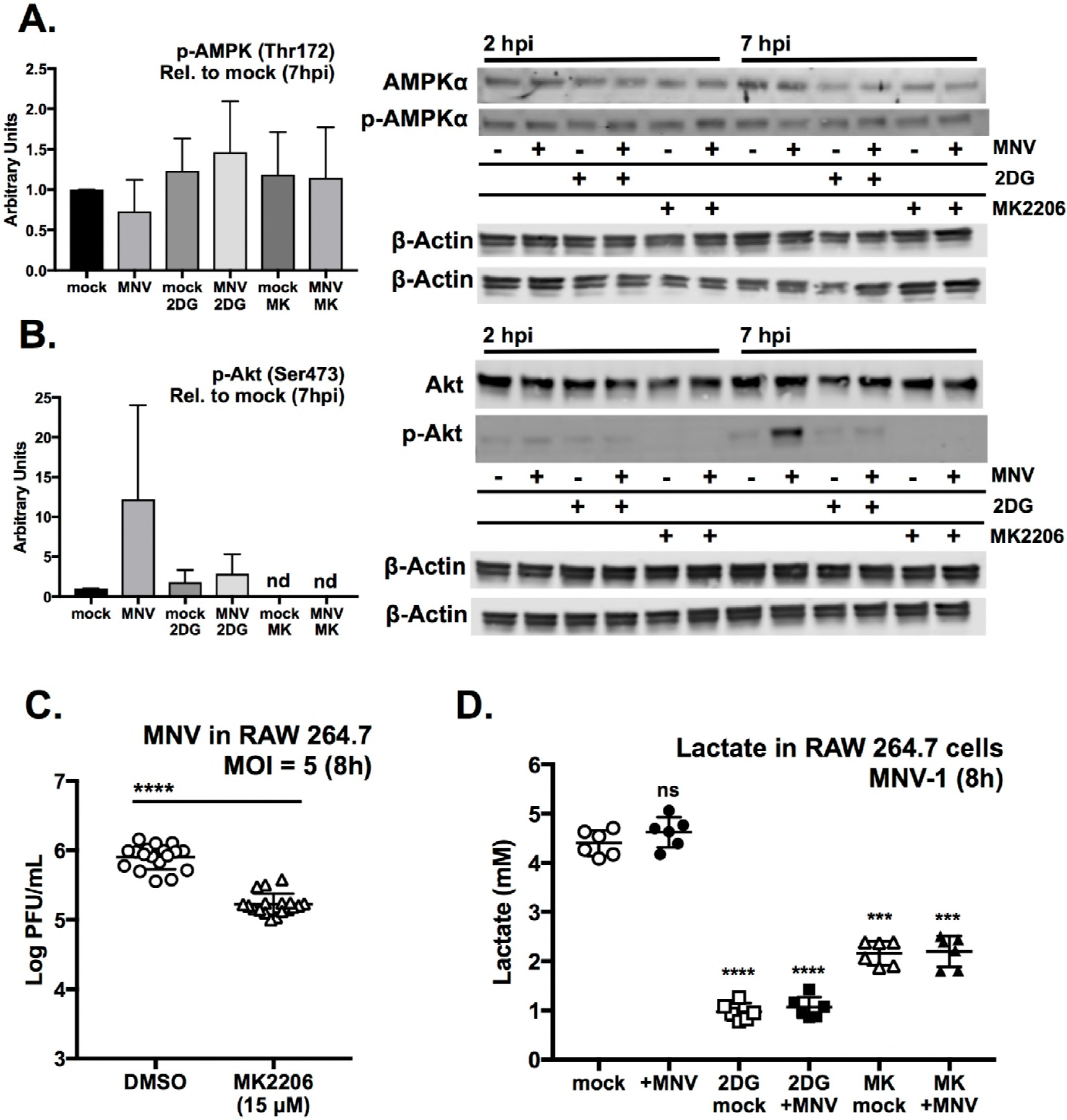
MNV upregulates glycolysis via Akt signaling. (A-B) Western blot analysis of RAW cells infected with MNV (MOI=5) for 2 and 7-hours for (A) AMPKα and phospho-AMPKα (Thr172), and (B) Akt and phospho-Akt (Ser473). Treatments were 10 mM 2DG and 15 μM MK2206. Western blot from 12hpi in FigS3. β-actin was used as loading control and for densitometry normalization. Graphs on the left represent densitometry analysis comparing protein phospho-protein relative to mock-infected cells at 7hpi. Graphs of densitometry analysis for 2hpi are in FigS4. (C) Inhibition of Akt phosphorylation with MK2206 reduces MNV-1 infection of RAW cells by about 0.75 log_10_. (D) Measurement of glycolysis via lactate production in mock and MNV-infected RAW cells after an 8-hour infection (MOI=5). Cells were treated with 10 mM 2DG and 15 μM MK2206. Mann-Whitney test used in where \*\*\*\**P*<0.0001 (combined three independent experiments). One-Way ANOVA used in with Dunnett’s multiple comparisons test (graph shows data for one of two independent experiments with three replicates each).

Another protein that has been implicated in energy sensing in multiple cell types is Akt. This kinase has been shown to play a key role in stimulating glycolysis and glucose metabolism via multiple mechanisms (72, 76, 84). In addition, Akt signaling is often altered during the infectious cycle of numerous viruses (85). Western blot analysis of Akt activation during MNV-1 infection demonstrated that Akt phosphorylation at Ser473 was slightly elevated at 2 hpi (~2-fold) (**Fig. 6B and Fig. S3**) above the baseline level of Akt activation in mock-treated RAW cells. Akt was further activated as indicated by the higher level of Ser473 phosphorylation at 7 hpi (~10-fold higher) (**Fig. 6B**) and 12 hpi (**Fig. S4**). 2DG treatment prevented these increases in Akt phosphorylation (**Fig. 6B**).

Because Akt phosphorylation was elevated during MNV infection and 2DG blocked Akt activation and viral replication, we asked whether inhibition of Akt signaling would inhibit MNV infection in RAW cells, linking Akt signaling with a change in host cell glycolysis. Treating cells with 15 μM MK2206, a potent inhibitor of Akt phosphorylation, completely prevented Akt phosphorylation at Ser473 (**Fig. 6B**) but did not affect AMPKα phosphorylation (**Fig. 6A**). Treating RAW cells with 15 μM MK2206 after the one hour MNV adsorption phase reduced viral production after eight hours by about one log_10_ (**Fig. 6C**). Furthermore, 2DG and MK2206 reduced RAW cell glycolysis irrespective of infection as measured by assaying end-point lactate production (**Fig. 6D**). Both compounds are non-toxic at these concentrations (**Fig. S1C and S1D**). These experiments demonstrate that Akt activation is a feature of MNV infection of RAW cells, and that Akt plays a role in maintaining glycolysis in these cells. Akt activation during MNV infection is consistent with a previous transcriptomic study of monocytes transfected with the non-structural protein NS1-2, which implicated NS1-2 in affecting PI3K-Akt signaling pathways (86). Taken together, these data are consistent with a model whereby MNV infection upregulates glycolysis via Akt signaling.

## DISCUSSION

When viruses infect cells, they are entirely dependent on the intracellular landscape of their hosts in order to replicate efficiently. Indeed the intracellular metabolic state of target cells acts as an intrinsic host factor, and a variety of metabolic pathways are important for successful viral infection (3). However, different viruses cause diverse metabolic effects in various cell types, and the mechanisms of viral engagement with host metabolic processes vary greatly (24–32). Thus, defining the specific host cell metabolic features that are required for individual viruses may reveal key host cell vulnerabilities that could be helpful for the future development of effective and safe antiviral therapies (46). Noroviruses lack effective therapies. In this study, we uncover central carbon metabolism as an intrinsic factor that is important for optimal infection of macrophages by MNV at early points during replication, suggesting a potential new anti-norovirus target.

Maintaining homeostasis of glucose metabolism in mammalian physiology is of importance in virtually every tissue, and glycolysis and OXPHOS are considered to be “central” carbon metabolism since they are a hub for multiple metabolic pathways, and their vital role in energy homeostasis. Therefore, it is not surprising that some viruses have evolved to take advantage of different aspects of these conserved pathways to their benefit. Interestingly, glycolysis may be increased or decreased in response to viral infection, with similar beneficial outcomes for the virus. For instance, dengue virus increases both glucose uptake and transcription of the important enzyme hexokinase 2 (28), while herpes simplex type 1 activates glycolysis by increasing transcription and activation of the enzyme phosphofructokinase-1 (PFK-1) (33), with both viruses relying on active glycolysis for optimal infection. On the other hand, Kaposi’s sarcoma-associated herpesvirus (KSHV) causes a suppression of both aerobic glycolysis and OXPHOS in transformed cells under nutrient stress, which thereby inhibits cell death and enhances viral survival in this model of the tumor microenvironment (24). Our observation that astrovirus infection was not affected by the treatment of Caco-2 cells with 2DG highlights that not all viruses require glycolysis in transformed cells, which generally conduct a significant level of “Warburg Effect” glycolysis at baseline (61). Thus, it was notable that 2DG inhibited MNV infection in non-transformed primary cells, highlighting the fact that carbon metabolism has pro-viral functions during norovirus infection. These results illustrate that the relationship of target cell metabolism to viral infection is cell type-specific and virus-specific.

Another notable aspect of the relationship between carbon metabolism and infection is the finding that glycolysis may facilitate infection outside of a canonical, metabolic role. HIV-1 causes an increase in expression of hexokinase-1 (HK1) accompanied by a decrease in enzymatic activity (87). Our findings with 2DG, which targets the enzymatic activity of hexokinase, points to a metabolic, rather than non-metabolic, role for glycolysis during norovirus infection. Specifically, glucose-6-phosphate (G6P), located at the intersection of glycolysis and PPP, is a major hub for macrophage metabolic regulation of MNV infection given that inhibition of the PPP also reduced viral infection.

One particular caveat of host cell metabolic profiling studies is the complexity of metabolic responses that immune cells can adopt in response to various stimuli. This is of particular relevance for macrophages. Although the M0/M1/M2 system of categorizing macrophage metabolic states is a useful construct for generalizing inflammatory versus non-inflammatory activity, these cells establish a complex range of metabolic phenotypes (22, 88, 89). For example, although the bacterial product LPS causes an increase in glycolysis and a decrease in OXPHOS in human monocytes, a different bacterial product, Pam3CysSK4 (P3C), causes both pathways to increase (23). Thus, two different bacterial products signaling through different Toll-like receptors (TLRs) establish unique metabolic profiles. This finding emphasizes that unique pathogens elicit complex host metabolic responses, and that the range of molecular signals that immune cells are responding to *in vivo* may determine the susceptibility of cell types to certain infections. The metabolomics survey in this study demonstrated that MNV infection, like P3C treatment, elicits an increase in both glycolysis and OXPHOS, and ongoing work is seeking to reveal the individual contributions of these two pathways to norovirus infection. In addition, since macrophages are target cells of MNV *in vivo* (44, 90), it is conceivable that their metabolic status during infection influences the establishment of norovirus infection at the cellular levels with potential influences on viral pathogenesis. However, future studies are needed to test this.

Another important aspect of macrophage metabolism is how metabolic rewiring controls functional outputs, such as microbial killing mechanisms and cytokine/chemokine production (7), which in turn could indirectly affect viral infection. Akt has been implicated in regulating reactive oxygen species (ROS) generation (75). Although MNV infection increased Akt activation, we did not observe an increase in general ROS in RAW cells upon 2DG treatment (**Fig. S5**). Furthermore, blocking glycolysis with 2DG did not cause a significant difference in the production of the inflammatory cytokine TNFα in 2DG-treated RAW cells during MNV infection (**Fig. S6**). Combined with the finding that the effect of 2DG is independent of type I IFN signaling, these data suggest that the antiviral effect of 2DG is not mediated via immune signaling. However, whether MNV affects general macrophage functions via Akt activation and metabolic rewiring of these cells will need to be tested in future studies.

A general caveat to the use of pharmacologic inhibitors in biological systems is their potential for inducing side effects. Although 2DG has been commonly used as a prototypical glycolysis inhibitor (59, 60), it may also affect other aspects of cell behavior that can influence infectivity. For example, 2DG has been shown to induce ROS-triggered autophagy via AMPK (91). This pathway is unlikely to be involved in the antiviral activity of 2DG in our studies considering the lack of ROS induction upon 2DG treatment in RAW cells (**Fig. S5**), and the lack of AMPK induction during infection. Another study showed that 2DG can be damaging for certain viral infections via initiation of an ER stress response in mice (92). Similarly, 2DG decreases porcine epidemic diarrhea virus infection *in vitro* via triggering the unfolded protein response and reducing protein translation (93). Our work showed that 2DG inhibited MNV infection early during the viral life cycle, affecting the translation of non-structural proteins and the transcription of new viral genomes. However, the virus does eventually begin to replicate genomes and produce viral proteins even in the presence of 2DG. Thus, the mechanism by which 2DG causes this lag in the MNV life cycle could be via a rapid cellular stress response, or a decrease in specific metabolites, or a combination of the two, and additional studies are needed to clarify the relative contribution of both.

Lastly, it should be noted that a variety of metabolites, including Ornithine, 3-Phospho-Serine, Creatinine among others, were also increased during MNV infection. While these molecules could be important host factors for viral infection, they were not explored further here. Such investigations and an extension of metabolic findings to human noroviruses are planned for the future. Human norovirus has remained stubbornly intractable to robust cultivation *in vitro*. Although there has been some success in infecting transformed B cells (40) and human intestinal enteroids (94) with human norovirus, viral loads remain low and an infectious, passagable cell culture-derived virus stock remains elusive (95). Identifying host cell factors such as metabolites and specific metabolic activities may therefore aid in optimizing *in vitro* cultivation systems for human noroviruses.

In conclusion, we have shown that central carbon metabolism in macrophages is an intrinsic factor promoting optimal infection of a norovirus. Our data are consistent with a model whereby MNV activates the protein kinase Akt to increase central carbon metabolism in macrophages. The glycolysis inhibitor 2DG inhibits norovirus (but not astrovirus) infection, independent of the type I IFN response by limiting non-structural protein translation and viral RNA synthesis. These findings reveal cellular metabolism as a potential therapeutic target for norovirus and suggest a new strategy for improving human norovirus culture systems.

## MATERIALS AND METHODS

Detailed methods can be found in Text S1 in the supplemental material.

### Compounds and reagents

Please refer to Text S1 in the supplemental material for details.

### Cell culture and virus strains

RAW 264.7 and Caco-2 cells were obtained from ATCC. The plaque purified MNV-1 clone (GV/MNV1/2002/USA) MNV-1.CW3 (43) (referred herein as MNV-1) was used at passage 6 in all experiments.

### Virus infections, virus transfection, and plaque assay

All MNV infections were done in the RAW 264.7 cell line, Balb/c primary bone marrow-derived macrophages (BMDM from male mice) or BMDM from WT and IFNAR1-knockout cells on a C57Bl6 background. Transfections and viral enumerations were performed similar to (62, 96, 97). Please refer to supplemental material for details.

### Cell viability assay

Cell viability was tested using Resazurin reagent according to the manufacturer’s recommendations (Biotium 30025-1).

### RNA extraction and RT-qPCR

Experiments were performed per manufacturer’s directions using Chloroform extraction (Trizol) or the Zymo Research Direct-zol RNA MiniPrep Plus (R2072).

### Strand-specific RT-qPCR

Strand-specific RT-qPCR for MNV was performed as previously described (63).

### Protein extraction, SDS-PAGE and immunoblotting

Experimental conditions and antibodies are detailed in supplemental material.

### Metabolomics assay

Samples were analyzed at the Michigan Regional Comprehensive Metabolomics Resource Core (MRC^2^) at the University of Michigan by Mass Spectrometry as detailed in supplemental material.

### Lactate assay

Cell supernatants were assessed for lactate using the Cayman Chemical Glycolysis Cell-Based Assay Kit (600450) per the manufacturer’s protocol.

### ELISA

Cytokine levels were determined at the University of Michigan Rogel Cancer Center Immunological Monitoring Core by ELISA (Duosets, R&D Systems, Minneapolis, MN) as detailed in supplemental materials.

### Statistical Analysis

Metabolomics data were analyzed in Metaboanalyst 4.0. For all other experiments, data were analyzed in Prism7 using tests as indicated in Figure legends.

## ACKNOWLEDGMENTS

This work was in part supported by NIH/NIAID R21/R33 AI102106 to C.E.W. and M.O.R. and the University of Michigan BMRC Bridging Support program. J.L. and I.G. are supported by grants from the Wellcome Trust (Ref: 207498/Z/17/Z) and the UK Biotechnology and Biological Sciences Research Council (Ref: BB/N001176/1). The work on this manuscript utilized Metabolomics Core Services supported by grant U24 DK097153 of NIH Common Funds Project to the University of Michigan. We thank Dr. Kim Green (NIH, NIAID, USA) for the ProPol antibody, Dr. Megan Baldridge (Washington University in St. Louis, MO, USA) for IFNAR−/− bone marrow, and the members of the O’Riordan and Wobus laboratories for suggestions.

## AUTHOR CONTRIBUTIONS

KDP, AOK, MXDOR, CEW conceived the experiments. KDP, AOK, JL, JRA, RJM carried out the experiments. KDP, AOK, JL analyzed the data. KDP, AOK, IG, MXDOR, CEW contributed to the interpretation of the results. KDP and CEW wrote the manuscript in consultation with IG and MXDOR.

